# First evaluation of a human DPP4 transgenic hamster model for MERS-CoV pathogenesis and transmission

**DOI:** 10.64898/2026.05.26.727115

**Authors:** Jacob Schön, Yanan Liu, Nico Joel Halwe, Tobias Britzke, Marie-Claire B. Codjia Risch, Rong Li, Nathan Merrill, Lorenz Ulrich, Jordi Rodon, Johanna Bork, Diana Bösel, Annika Beyer, Marcel A. Müller, Christian Drosten, Angele Breithaupt, Donata Hoffmann, Zhongde Wang, Martin Beer

## Abstract

MERS-CoV poses a constant pandemic risk, as its viral lineages continue evolving, and zoonotic spillover events could lead to random viral polymorphisms that might lead to human adapted variants. Currently, no small animal model reliably recapitulates both disease progression and transmission dynamics, which are critical aspects for counter-viral measures like vaccine development. Although the Syrian hamster is an optimal animal model for SARS-CoV-2 infection and transmission, it is naturally resistant to MERS-CoV infection. Dipeptidyl peptidase-4 (DPP4) is the functional receptor for MERS-CoV infection, and is highly expressed in human kidney, intestine, liver, and lung tissues. Here, we evaluated the suitability of a human DPP4 (hDPP4) transgenic Syrian hamster model for MERS-CoV research. We used two different MERS-CoV strains (EMC/2012 and D10540/2023) for intranasal inoculation of hamsters. Both strains replicated efficiently, led to comparable severe clinical outcomes, and had similar viral transmission efficiencies. MERS-CoV RNA and nucleoprotein antigen were mainly detected in the brain and the respiratory tract. In summary, we validated a novel hDPP4-transgenic hamster as a suitable model for MERS-CoV infection enabling vaccine and transmission research.

## Introduction

The Middle East respiratory syndrome coronavirus (MERS-CoV) is a zoonotic respiratory *Betacoronavirus* in the family of *Coronaviridae*. Along with the *Alpha-* and *Betacoronaviruses* HKU1, 229E, NL63, OC43, SARS-CoV and SARS-CoV-2, MERS-CoV belongs to one out of seven currently known coronaviruses capable of infecting humans ^1^. Initially, MERS-CoV was identified and isolated in 2012 from a 60-year-old male patient in Saudi Arabia suffering from acute pneumonia and renal failure with a fatal outcome ^2^. Until May 2025, over 2,637 human infections with MERS-CoV from 27 countries had been laboratory-confirmed, with a case-fatality ratio close to 36% ^3,4^. The clinical picture can also include fever, coughing and shortness of breath, as well as gastrointestinal symptoms ^3–5^.

The virus is naturally circulating in dromedary camels, which are therefore considered the primary reservoir of the virus ^6–8^. Camel-to-human interactions lead to frequent spillover infections into human population, with subsequent limited human-to-human transmission

Hereafter, strong camel-to-human interfaces lead to frequent spill-over infections into the human population with subsequent human-to-human transmission also occurs to some extend ^9–11^. Phylogenetic analysis divides MERS-CoV into three independent clades, clade A, B and C^12,13^. Clade A MERS-CoV was responsible for the first human respiratory disease outbreak in 2012 in the Arabian Peninsula, but has been replaced by clade B MERS-CoV strains ^14–16^. Clade B strains are further subdivided into seven subclades, of which subclade B5 predominated since 2015 and has caused the majority of human outbreaks ^12,14,15^. Nevertheless, the currently circulating clade B5 MERS-CoV has not yet been comprehensively characterized, particularly *in vivo*. Clade C MERS-CoV circulation appears to be limited to dromedary camels in African countries, causing only sporadic and mostly asymptomatic zoonotic infections and no associated outbreaks ^13,14,17–22^. Self-limiting human-to-human transmission has been confirmed, especially through direct contact with infected individuals, including household contacts or healthcare workers in a nosocomial context ^23–25^. Even though the numbers of human cases have declined since the SARS-CoV-2 pandemic, ongoing spillover events into human populations pose a public health threat with pandemic potential ^26^.

There are a limited number of animal models to study MERS-CoV. While ferrets and Syrian hamsters stood out as excellent animal models for other viral respiratory pathogens, such as Influenza A or SARS-CoV-2 infection, these animals are naturally not susceptible to MERS-CoV infection ^27–30^. MERS-CoV enters its host cells by binding to the cell surface protein Dipeptidyl peptidase-4 (DPP4) ^31^. It is believed that ferret DPP4 and hamster DPP4 do not support MERS-CoV attachment ^30^, in contrast to the human DPP4 (hDPP4). To address this, we generated a hDPP4 transgenic Syrian hamster model using a PiggyBac-mediated transgenic approach with pronuclear injection of this vector into hamster zygotes ^32–34^. In this model, hDPP4 expression is driven by the human cytokeratin 18 promoter. This transgenic hamster line showed normal development and fertility and was used in the subsequent experiment. Additional details on the generation of the hDPP4 hamsters are provided in Wang et al., 2026.

Here we evaluated novel hDPP4 Syrian hamster as a small rodent animal model to study MERS-CoV infection, with a focus on pathogenesis and transmission. We intranasally inoculated the hamster model with the prototypic MERS-CoV EMC/2012 (clade A) and the more recent virus strain 10540/2023 (clade B5) circulating in dromedary camels ^9^. Both isolates efficiently replicate and cause severe disease in the hDPP4 hamster model. Importantly, we also observed efficient MERS-CoV transmission among animals in direct contact. Our findings demonstrate that hDPP4 hamsters serve as a reliable *in vivo* model for investigating MERS-CoV pathogenesis and transmission, which further support their use in preclinical evaluation of countermeasures, such as antivirals and vaccines.

## Results

### hDPP4 Syrian hamsters are susceptible to MERS-CoV infection and develop fatal disease

We intranasally inoculated transgenic hDPP4 Syrian hamsters with 10^5.5^ TCID_50_ per animal of either MERS-CoV EMC/2012 (Group 1, “EMC-Group”; n = 7) or with MERS-CoV 10540/2023 in the second group (“10540-Group”; n = 7) (**Figure 1a**). Each animal was caged individually, and one day post infection (dpi) a non-infected naïve direct contact animal was co-housed to an inoculated donor animal, forming five transmission pairs per group (**Figure 1a**). Two inoculated (donor) animals per group were sacrificed at 4 dpi to evaluate viral organ dissemination during the suspected acute infection phase (**Figure 1a**). All inoculated animals, regardless of the used virus isolate, developed severe clinical signs, including pronounced head tilt, marked neurological symptoms, and an unsteady ataxic gait (**Supplementary Table 1**). One inoculated animal in the EMC group succumbed to disease without showing clinical signs before, while all remaining animals in both groups reached humane endpoint criteria and were euthanized between 5 and 7 dpi (**Figure 1b**). Body weight loss started abruptly at 3 dpi, and already by 5 dpi, two directly inoculated hDPP4 hamsters (one per group) were euthanized due to reaching humane endpoint criteria based on clinical signs (**Figures 1b and c, Supplementary Table 1**). All remaining donor hamsters in both groups were euthanized at 6 or 7 dpi upon reaching humane endpoint criteria, with ataxia being the dominant clinical sign, indicating involvement of the central nervous system (**Supplementary Table 1**). MERS-CoV RNA was detected by RT-qPCR analysis in nasal wash samples throughout the study, indicating efficient virus replication and shedding in all inoculated animals of both groups from 1 dpi onwards (**Figures 1d and e**).

**Figure 1:**
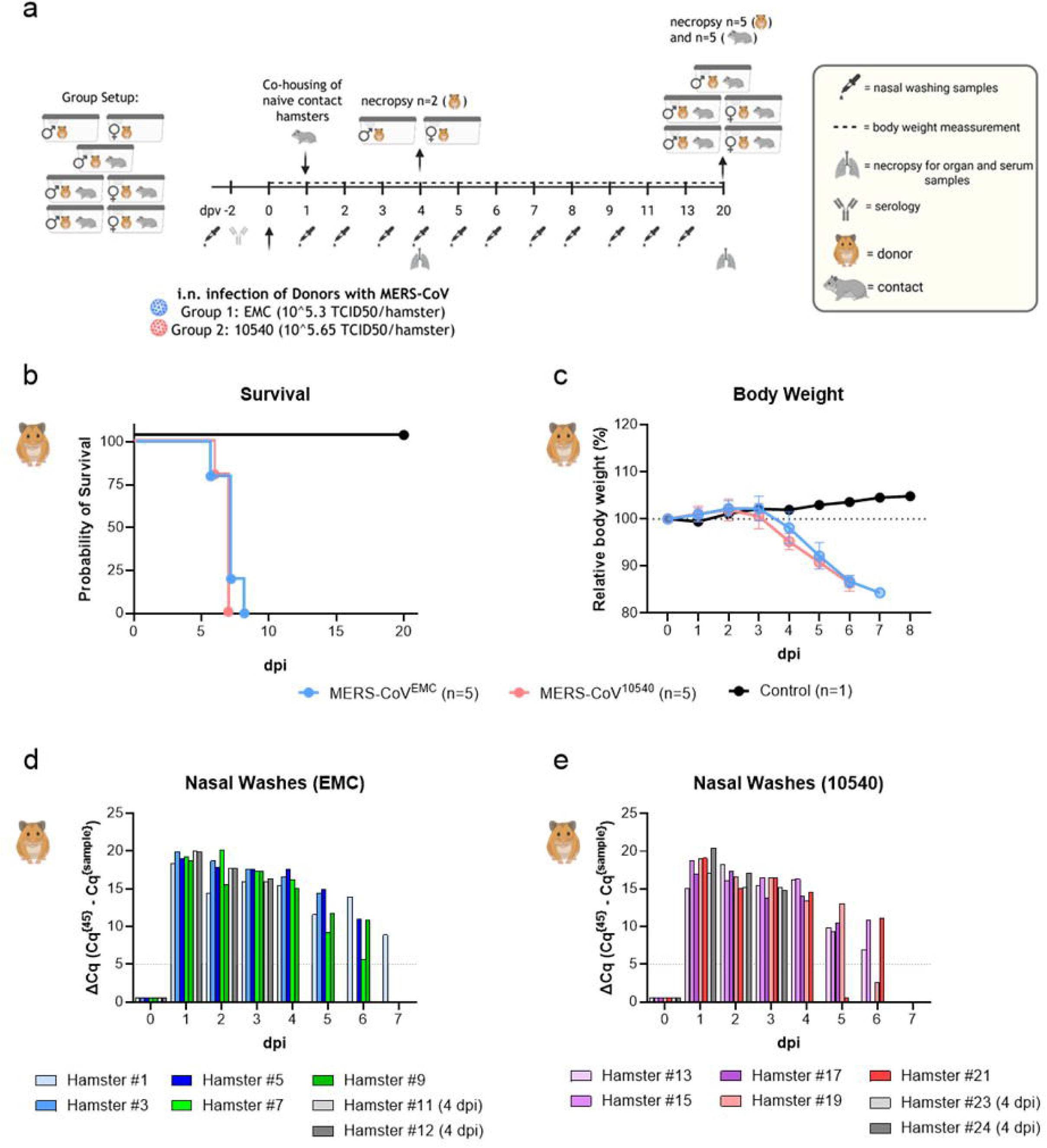
Inoculation of hDPP4-Syrian hamsters with two different MERS-CoV strains. (a) Experimental design: Male and female animals were intranasally inoculated (donor animals) with either the EMC/2012 or the 10540/2023 MERS-CoV strains. One day post inoculation (1 dpi), naïve direct-contact animals were co-housed with donors to assess transmission. (b) Survival of donor animals; one EMC-donor animal succumbed to disease, and the remaining EMC- and 10540-donor animals were euthanized upon reaching human end-point criteria. (c) Relative body weight (%) of donor animals over time following MERS-CoV inoculation. (d) Nasal wash samples from the EMC group and (e) from the 10540 group were analyzed by RT-qPCR for detection of MERS-CoV genome. Assay background limit is indicated with a dotted line.

### Efficient MERS-CoV transmission to direct-contact animals

At 1 dpi, naïve direct-contact animals (n = 5 per group) were co-housed with infected donors in a 1:1 setup per cage ((males (n=15) and females (n=10)) with same sexes caged together (**Supplement Table 1**)), resulting in five transmission pairs per group. Contact animals in both groups developed clinical signs similar to those observed in donor animals and reached the humane endpoint criteria between 8 and 12 dpi (**Figure 2a, Supplementary Table 1)**. One contact animal in the EMC group succumbed to disease at 9 dpi (**Figure 2a**). Overall, one out of five EMC-contact animals and two out of five 10540-contact animals survived until the end of the experiment at 20 dpi without developing obvious clinical signs (**Figure 2a**). Nevertheless, this was accompanied by body weight loss starting from 5 dpi (**Figure 2b**). To further assess transmission efficacy, nasal wash samples of contacts were analyzed for MERS-CoV RNA presence. All contact animals in the EMC group tested positive for viral genomes by 6 days after contact (7 dpi) at latest (**Figure 2c**). A similar pattern was observed in the 10540 group, although one contact animal tested positive later by 8 dpi (**Figure 2d**). Notably, some Cq values were high and only intermittently positive at individual time points. For example, in transmission pair 5 (hamster #19 and #20) of the 10540 group, viral genome was detected only sporadically with Cq-values of 37.1 (3 dpi), 34.5 (5 dpi), and 36.3 (6 dpi).

**Figure 2:**
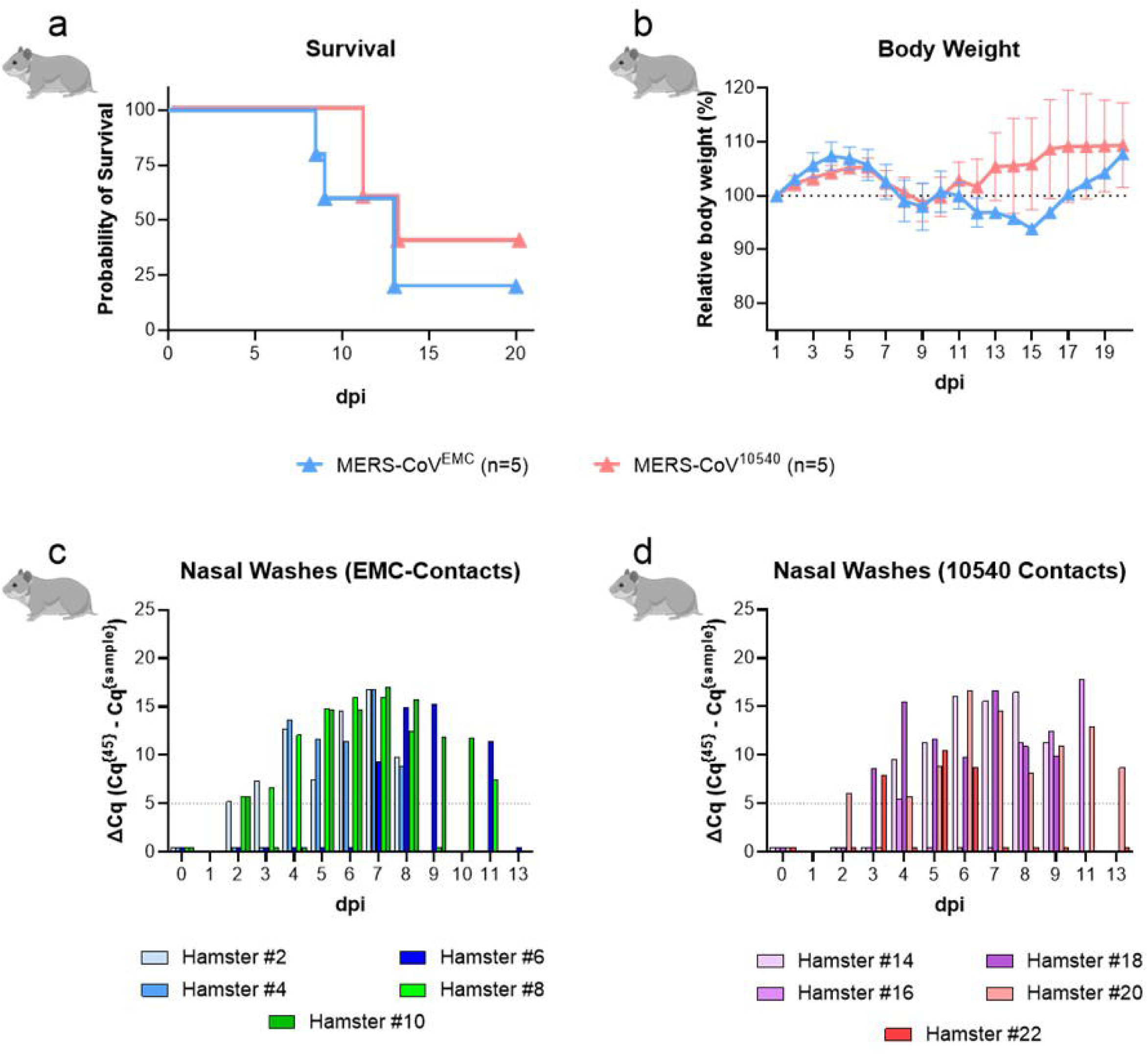
Virus transmission from MERS-CoV inoculated hDPP4 Syrian hamsters to naïve direct-contact hamsters. Five donor animals were co-housed with naïve direct-contact animals 1 day post inoculation (dpi) in a 1:1 setup per cage, forming five transmission pairs for each group (EMC or 10540). (a) Survival of naïve direct-contact hDPP4 hamster, following co-housing to MERS-CoV-inoculated donor hamsters. (b) Relative body weight (%) of contact animals over time following co-housing to donor animals. (c) Nasal wash samples from contact animals following co-housing to EMC-donor animals or to (d) 10540-donor animals were analyzed by MERS-CoV RT-qPCR.

### MERS-CoV disseminates primarily in nose, lung, and brain

At 4 dpi, two intranasally inoculated animals per MERS-CoV strain were sacrificed to analyze a broad panel of tissues for MERS-CoV presence. The same panel was assessed for all the later sacrificed animals, as most had to be euthanized around 6 dpi. All donor animals, regardless of the isolate, showed high viral RNA loads between 4 and 7 dpi in the central nervous system (CNS) samples (Cerebrum and Cerebellum), as well as in samples from lung and nasal turbinates to a lower extent (**Figure 3a**). Especially early time point samples also tested positive for infectious virus by endpoint titration in Vero E6 cells (**Supplementary Figure 1**). All other samples tested negative by RT-qPCR or only weakly positive including heart, liver, kidney and spleen, as well as rectal swabs (**Figure 3a**). Of note, one animal of the EMC-Group died prematurely at 7 dpi (animal #1) and was tested highly positive in the kidneys (**Figure 3a**). Neutralizing seroresponses in inoculated donor animals could be detected as early as 6 dpi (**Figure 3b**). Nevertheless, the immune responses in the donor animals were not sufficient to protect from severe clinical disease. Interestingly, the neutralizing titers in the EMC group were higher than those in the 10540 group, with no major differences if EMC or 10540 was used as test virus (**Figure 3b**). A similar viral distribution pattern was observed for the direct contact animals of both groups (**Figure 3c**). The one EMC-contact animal (animal #6) and the two 10540-contact animals (animal #20 and #22), which survived until the end of the observation period at 20 dpi had only residual traces of viral genomes in the CNS or the lung (animal #6 and #20) (**Figure 3c**), while one contact animal (#22) of the 10540 group was never productively infected; hence all tissue samples tested negative MERS-CoV RNA and it did not seroconvert (**Figure 3d**).

**Figure 3:**
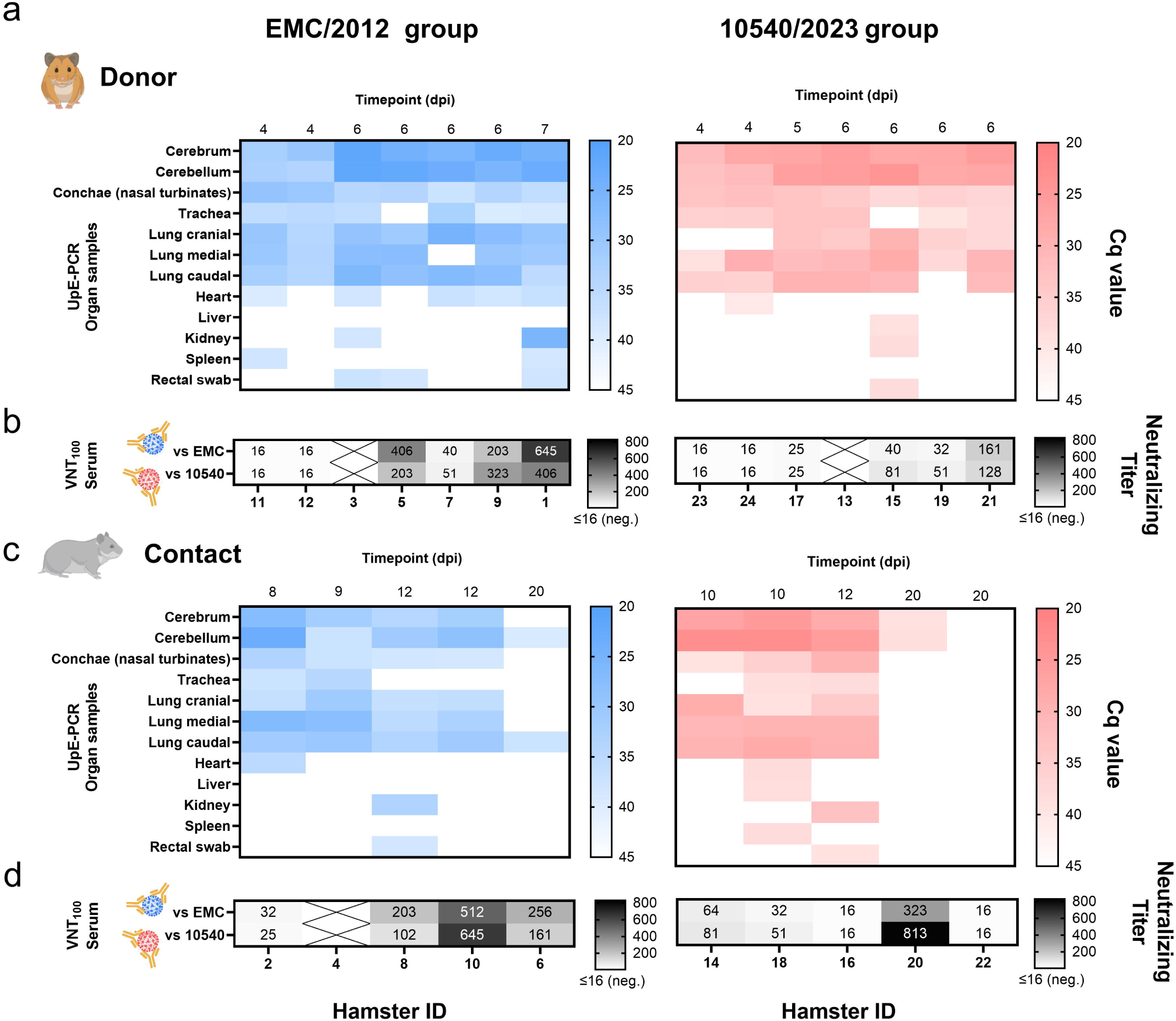
MERS-CoV genome detection across multiple organs and corresponding neutralizing humoral immune response. (a) Hamsters inoculated with MERS-CoV EMC or 10540 strains exhibited high viral genome loads in the central nervous system (CNS) and in the upper and lower respiratory tract tissues between 4 and 7 dpi. (b) Virus-neutralizing titers were assessed in serum samples by neutralization assay. (c) Viral RNA detection in a broad panel of organs from direct-contact animals exposed to EMC- or 10540-donor hamsters. (d) Neutralizing antibody responses in serum samples from direct-contact animals, shown as virus-neutralizing titers. The limit of detection (LOD) was defined as the starting serum dilution of 1:16; samples with no detectable neutralization were assigned a value of <16.

### Cellular localization of MERS-CoV in infected tissues

We next identified target cells in tissues and organs with high viral genome loads at 4 dpi and also on 20 dpi using immunohistochemistry to stain the nucleoprotein of MERS-CoV. Consistent with the MERS-CoV RNA viral load data of donor animals, both virus variants exhibited similar antigen pattern. Viral antigen was detected in multiple cell types within the nasal mucosa: in olfactory sensory neurons (**Figure 4a**), in sustentacular cells (**Figure 4b**, arrow), in ciliated epithelial cells of the respiratory epithelium (**Figure 4c**), and in Bowman’s glands (**Figure 4d**, arrow). Furthermore, antigen was found in peripheral nerves (**Figure 4e**, arrow), indicating involvement of neuronal structures beyond the sensory epithelium. In the lung, we detected virus protein in the bronchial epithelium, including ciliated epithelial cells (**Figure 4f**), as well as in the alveolar epithelium (**Figure 4g**), while in the brain virus protein was prominent in neurons (**Figure 4h**, arrow) and some glial cells (**Figure 4i**, arrow). No antigen was detected on animals euthanized at 20 dpi, suggesting viral clearance.

**Figure 4:**
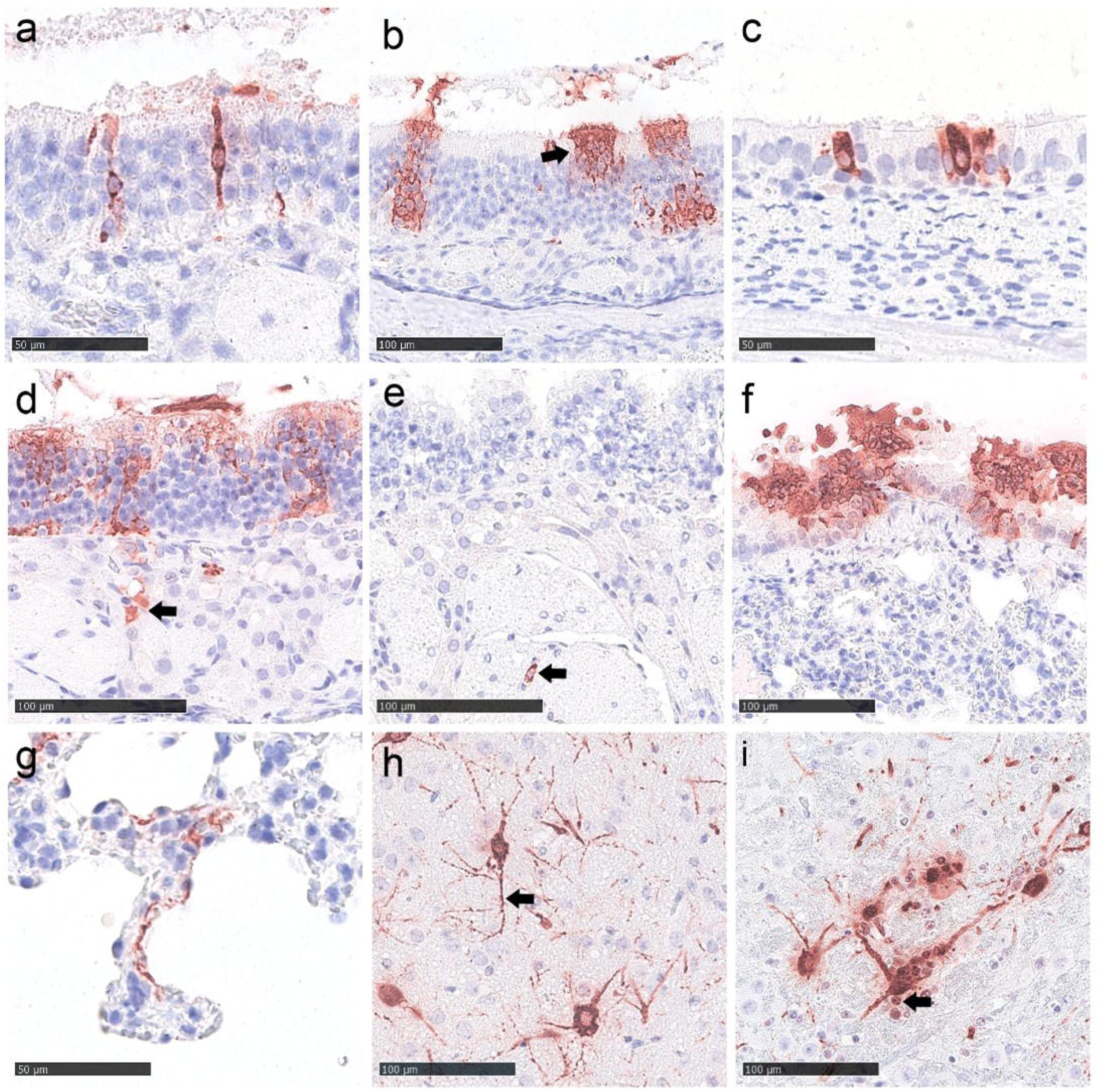
MERS-CoV antigen detection in tissues of the hDPP4 Syrian hamster model at 4 days post infection. (a–e) Nasal cavity with MERS-CoV antigen detection in olfactory sensory neurons (a), sustentacular cells (b, arrow), ciliated respiratory epithelial cells (c), Bowman’s glands (d, arrow), and peripheral nerves (e, arrow). (f, g) Lung showing MERS-CoV antigen in ciliated bronchial epithelium (f), and alveolar epithelium (g). (h, i) Brain with MERS-CoV-protein detection in neurons showing prominent labeling of neuronal processes (h, arrow), and in glial cells (i, arrow). Immunohistochemistry was performed using an anti-MERS-CoV nucleoprotein antibody. Red-brown signal was visualized using 3-amino-9-ethylcarbazole (AEC), with hematoxylin counterstaining. Scale bars: 50 µm (a, c, g); 100 µm (b, d–f, h, i).

### Human DPP4 expression in tissues is comparable to that of endogenous hamster DPP4 expression but it does not fully correlate with virus dissemination

To evaluate specific expression of the human DPP4 transgene or the endogenous hamster DPP4 in tissues, we compared mRNA levels of both DPP4 orthologs by RT-qPCR relative to Peptidylprolyl isomerase A (PPIA) housekeeping gene mRNA levels, with experimentally validated species-specific RT-qPCR primers for hDPP4 and hamster DPP4, respectively (**Supplementary Figure 2**). Primer and probe sequences are listed in Table 1. It appears that for the tested tissues the mRNA levels are mostly at a similar level, except for heart tissue, in which the hDPP4 was more than ten-fold higher expressed than the hamster counterpart (**Figure 5a**). To evaluate this also at protein level, we tested brain, lung, trachea, heart, kidney and liver of hDPP4 hamster in comparison to its counterpart in wild type by Western blotting for DPP4 protein abundance. Similar to the results from RT-qPCR, Western blotting showed high hDPP4 expression in lung, heart, kidney, and liver, whereas expression was barely detectable in brain, trachea, and spleen (**Figure 5b**); considering the relative low levels of expression of hDPP4 detected by RT-qPCR in brain, trachea, and spleen, it is most likely that the low sensitivity of the anti-hDPP4 antibody failed to detect hDPP4 expression with Western blotting. Signals were also detected in some wild type hamster tissues, including lung and kidney, where RT-qPCR showed the highest levels of hamster DPP4 expression, indicating that the anti-hDPP4 antibody is also weakly cross-reactive with hamster DPP4; thus, the protein levels detected by Western blotting reflects the cumulative abundance of hDPP4 and hamster DPP4 (**Figure 5b**).

**Figure 5.**
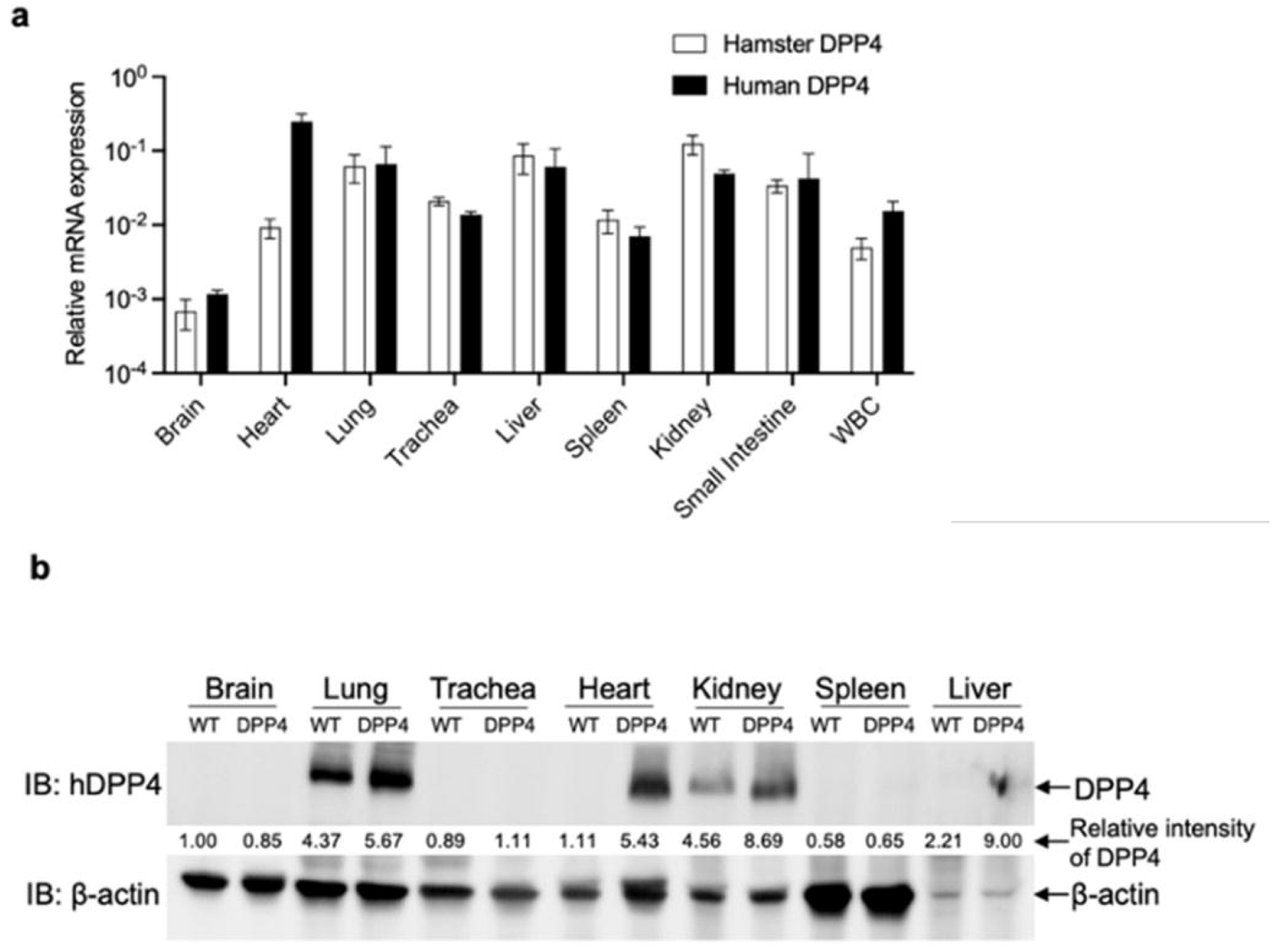
Characterization of hDPP4 expression across different organs in hDPP4 transgenic hamsters. (a) RT–qPCR analysis of hDPP4 expression in comparison to hamster DPP4 in different tissues. Total RNA was extracted from the indicated tissues and subjected to ortholog-specific RT–qPCR analysis. Hamster PPIA gene was used as the internal control. Relative expression of hDPP4 and hamster DPP4 was calculated by normalization to hamster PPIA. Data are shown as mean ± SD from three animals. (b) Detection of hDPP4 by Western blotting in different tissues from wild-type and hDPP4 hamsters. The relative intensity of hDPP4 and hamster DPP4 was normalized to β-actin in the brain tissue of a WT hamster.

**Table 1:**
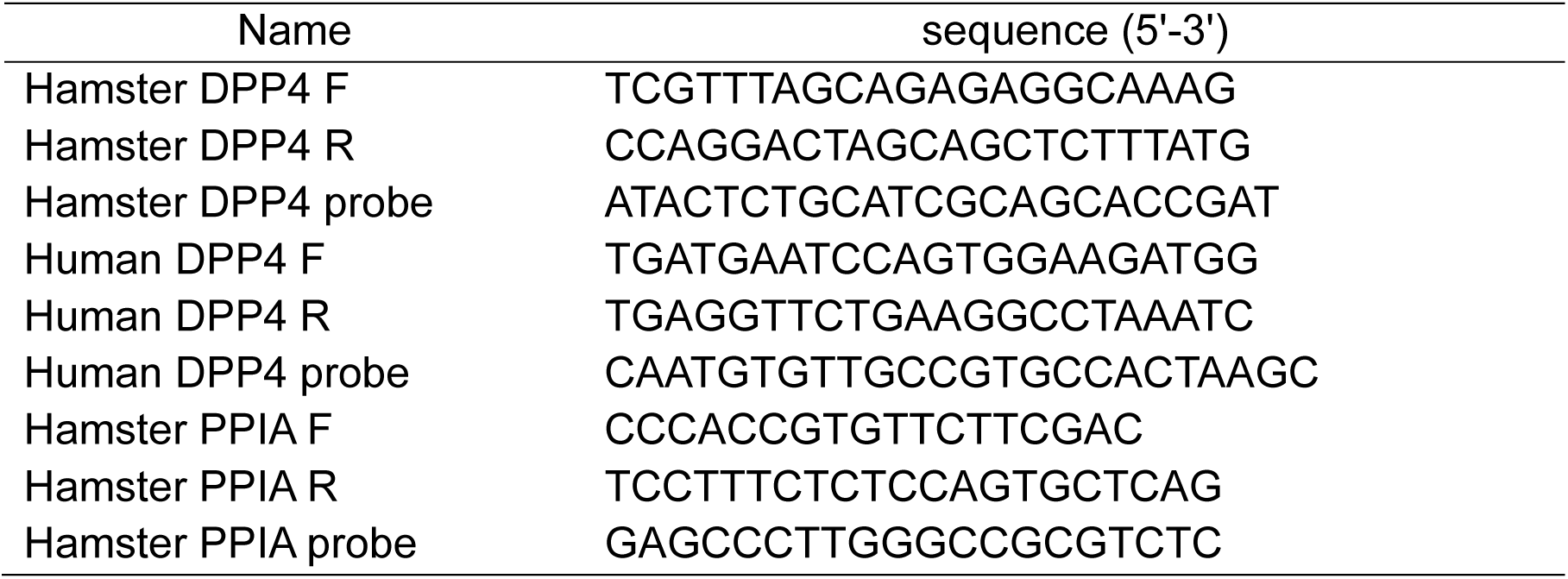
Primers and probes for hamster *DPP4*.

## Discussion

With an estimated case-fatality rate of 36% and frequently occurring spillover infections from camels to humans, MERS-CoV poses a constant pandemic threat. To better understand pathogenesis and to support the development of both prevention and treatment strategies against MERS-CoV, small animal models that recapitulate disease and transmission in humans are crucial. As MERS-CoV does not naturally bind to the mice or hamster DPP4 receptor, wild type animals cannot be readily used for these studies ^35^. Several attempts were made to generate small animal models for MERS-CoV, including adenovirus (Ad5) mediated introduction of hDPP4 into mouse lungs ^36^, global transgenic expression of hDPP4 under a CAG promotor in mice, ^37^ as well as using the keratin 18 (K18) promotor for epithelium specific expression ^38^. While the adenovirus humanized mice developed an interstitial pneumonia with no fatal outcome ^36^, the CAG- ^39^ and the K18-hDPP4 mice developed severe clinics including infection of the CNS and fatal outcome ^38,40^, which makes them favorable for vaccine testing due to the unambiguous readout ^41^. Nevertheless, none of these models demonstrated sufficient viral transmission to contact animals ^38,40^, rendering them unsuitable for transmission studies or for evaluating the potential of vaccines to reduce transmission and induce “sterile immunity”.

A robust transmission model is therefore still lacking. For other respiratory viruses, such as SARS-CoV-2, the Syrian hamster has proven to be a highly suitable and valuable model for pathogenesis and transmission studies. To enable the use of this otherwise promising model, we generated the human DPP4 transgenic hamster model. The generation of the hDPP4 transgenic hamster model is described in detail by Wang et al. (2026).

Here, we further evaluated this novel small animal model for its susceptibility to MERS-CoV by using two different lineages: the EMC reference virus from 2012 and a recently circulating B5 lineage strain (10540) from 2023 of camel origin ^9^. Our experiments clearly demonstrated that all inoculated animals have been productively infected and developed severe disease, reaching humane end-point criteria within a week post infection. Histopathological and immunohistochemical analyses confirmed infection of multiple respiratory epithelial cell types, reflecting key aspects of airway and lung involvement. However, prominent infection of the CNS was also observed, suggesting that severe disease progression is associated with viral replication and immune responses within the CNS. Although this does not fully reflect the natural pathogenesis in humans, which is primarily characterized by pneumonia ^42^, the consistent and well-defined clinical outcome may represent an advantage for vaccine efficacy studies.

Comparing the tissue distribution of MERS-CoV (Figure 3a, c) with hDPP4 transgene expression revealed that, except for the lungs—where high hDPP4 mRNA and protein abundance correlated with high viral genome loads—the relative levels of hDPP4 expression (Figure 5a) did not directly correlate with the amounts of MERS-CoV RNA detected in the tissues (Figure 3a). For example, despite relatively low hDPP4 mRNA and protein expression in the brain, high levels of MERS-CoV RNA were detected, suggesting either that even low hDPP4 expression may be sufficient to support MERS-CoV replication and dissemination, or that alternative tissue-specific pathways may contribute to viral spread. Overall, MERS-CoV RNA levels correlated more closely with DPP4 protein expression than with hDPP4-mRNA expression. No viral RNA was detected in heart and liver tissues, which also exhibited low DPP4 protein levels. In contrast, the lungs showed both high DPP4 protein abundance and high viral RNA loads. However, this correlation did not hold true for the kidneys, where high levels of DPP4 protein were detected despite absent or very low viral RNA levels (with the exception of kidney tissue from animal #1 of the EMC group; Figure 3a). Taken together, these data indicate that hDPP4 protein abundance alone does not fully explain MERS-CoV genomic loads across tissues, suggesting that additional mechanisms of viral entry or dissemination (under certain conditions, a partially DPP4-independent or auxiliary attachment factors or co-receptors involving replication) may operate once infection is established. Nevertheless, technical limitations, such as low sensitivity for DPP4 detection in certain tissues, should also be considered and will require further investigation. In addition, hDPP4 reactivity was also detected in wild-type hamster lungs, indicating a degree of antibody cross-reactivity and limited species specificity of the antibody used.

Importantly, both isolates transmitted efficiently to direct-contact hamsters, with transmission rates of 80% (4/5) for strain 10540 and 100% (5/5) for the EMC strain. Disease progression in the contact animals was comparable to that observed in the donor hamsters. These findings are consistent with the data reported by Wang et al. (2026) and address a major gap in MERS-CoV research that has remained unresolved due to the limitations of currently available small animal models.

These characteristics make the hDPP4 hamster model particularly valuable for investigating the determinants of viral transmission and for evaluating intervention strategies, such as antivirals and vaccines, aimed not only at reducing disease severity but also at preventing MERS-CoV shedding and transmission. In contrast to existing hDPP4 mouse models ^38,40^, hDPP4 hamsters support not only MERS-CoV replication in the respiratory tract and CNS, but also efficient inter-animal transmission. As a result, transgenic hDPP4 hamsters may become a highly useful model for MERS-CoV research, as they can be maintained in most laboratory settings while better accommodating scalability and biosafety requirements. Their significance for MERS-CoV studies may ultimately parallel the role of hACE2 transgenic hamsters in SARS-CoV-2 research ^43–47^. However, the precise mode of MERS-CoV transmission between donor and contact hamsters remains to be determined in future studies, including the relative contributions of direct contact, fomite transmission, and airborne transmission over longer distances.

A key limitation of this model is highlighted by our observation of pronounced neurotropism for both MERS-CoV strains in hDPP4 hamsters, a feature that does not fully recapitulate the pathology observed in severe human MERS cases. Nevertheless, compared with previously established models, including hDPP4-transgenic mice and non-human primates, the hamster model combines scalability with functional virus transmission. Similar to other animal models, however, this system also involves trade-offs, including species-specific differences in immune responses and a more severe disease phenotype than typically observed in humans. In this regard, well-established hACE2-transgenic mouse models of SARS-CoV-2 have likewise demonstrated substantial experimental utility despite exhibiting additional features such as marked neurotropism ^48^. Furthermore, the pronounced disease severity of these models represents an important advantage for evaluating intervention strategies ^49^.

Although MERS-CoV spike protein activation is thought to occur predominantly through intracellular cleavage by furin or related proteases, it remains possible that a subset of virions is released in an incompletely processed state ^50^. Such particles could subsequently depend on tissue-specific expression levels of proteases for activation, including type II transmembrane serine proteases (TTSPs) or cathepsins ^50^. Together with the presence of the receptor the tissue-specific expression of these host proteases may therefore contribute to viral tropism and pathogenicity. Further *in vitro* and *in vivo* studies will be required to address these questions, as they were beyond the scope of the present initial *in vivo* characterization study.

In summary, we established a preclinical small-animal model that supports efficient MERS-CoV infection and transmission. Together with the parallel study by Wang et al., our findings provide complementary and expanded characterization of this model, strengthening the overall evidence for its relevance and utility. The detection of viral antigen in the lungs recapitulates key aspects of human MERS pathogenesis, highlighting the model’s value for translational research, despite the fact that this is a lethal model of MERS-CoV infection suffering from acute encephalitis.

Importantly, this model may also provide a unique platform for studying cross-protective immunity between SARS-CoV-2 and MERS-CoV. Most currently available small-animal models are permissive to either SARS-CoV-2 or MERS-CoV infection, but not both, thereby limiting direct investigation of heterologous coronavirus immunity. In contrast, our model enables studies assessing whether prior SARS-CoV-2 infection or vaccine-induced immunity modulates disease outcome following subsequent MERS-CoV challenge. Such investigations may help clarify whether the marked decline in reported MERS cases following the COVID-19 pandemic was influenced, at least in part, by cross-reactive immune responses induced by widespread SARS-CoV-2 exposure.

Future studies should further elucidate the spatiotemporal interplay between viral tropism, hDPP4 expression, host protease distribution, and tissue injury.

## Methods

### Viruses

Two wild-type recombinant MERS-CoV strains, EMC/2012 (clade A; GenBank Accession No. JX869059.2) and KSA/F6-P4a/D10540/2023 (clade B lineage 5; GenBank Accession No. PP952189.1), were rescued by previously established reverse genetics systems ^51,52^. The genomic sequence of the MERS-CoV 10540 strain was obtained in a previous study ^9^. We completed its full-genome sequence by 5’ and 3’ RACE PCR, using the Template Switching RT Enzyme Mix (New England Biolabs) and the PrimeScript RT Reagent Kit (Takara) oligo(dT) method to generate viral cDNA according to the manufacturer’s instructions, respectively. Viral genome-end cDNA was amplified with the KOD Hot Start Master Mix (Sigma-Aldrich) and sequenced via Illumina shot-gun technology (REF No. 10) and Sanger sequencing. The transformation-associated recombination approach was used to clone the full genome into yeast-artificial chromosome and rescue a functional wild-type 10540 infectious virus. Briefly, the full-genome of MERS-CoV 10540 strain was amplified from viral RNA via twelve conventional RT-PCR assays. PCR products were assembled into a pCC1-His vector using highly transformable *S. cerevisiae* VL6-48N and screened for their correct assembly. The resulting plasmids were propagated in *E. cloni* 10G bacteria, isolated, linearized, and subjected to *in-vitro* transcription. The viral RNA was transfected into BHK-J cells and the recombinant MERS-CoV 10540 strain was eventually rescued, propagated in Vero E6-TMPRSS2 cells, and purified for subsequent functional assays. Additional data regarding the design and performance of the reverse genetics system is in preparation by Rodon et al.

### Cells

VeroE6 cells (Collection of Cell Lines in Veterinary Medicine CCLV-RIE 0929) were cultured using a mixture of equal volumes of Eagle MEM (Hanks’ balanced salts solution) and Eagle MEM (Earle’s balanced salts solution) supplemented with 2 mM L-Glutamine, nonessential amino acids adjusted to 850 mg/L, NaHCO_3_, 120 mg/L sodium pyruvate, 10% fetal bovine serum (FBS), pH 7.2.

### Animal ethics and declarations

The hDPP4-hamster experiment was evaluated by the responsible ethics committee of the State Office of Agriculture, Food Safety, and Fishery in Mecklenburg–Western Pomerania (LALLF M-V), and gained governmental approval under registration number LVL MV TSD/7221.3-1-059/24.

### hDPP4-hamster study

Twenty-five transgenic golden Syrian hamsters (15 males, 10 females, age of 12 weeks) expressing the human DPP4 receptor (hDPP4-hamster) in addition to the hamster DPP4-receptor were shipped to the Friedrich-Loeffler-Institute from the “Laboratory Animal Research Center” at the Utah State University, USA. A detailed experimental setup of the animal trial is depicted in **Figure 1a**. Twelve hamsters each were allocated into two different experimental groups (group 1-MERS-CoV^EMC^, group 2-MERS-CoV^10540^) comprised of in total six cages (two animals per cage). One additional hamster served as uninfected negative control. The first five cages of each group contained one intranasally inoculated donor hamster, as well as one uninfected contact hamster. The contact hamsters were strictly separated from the donor hamsters prior to inoculation, and were added back into the cage with the appropriate donor hamster at one day post infection (1 dpi) to prevent unspecific transmission from the inoculum reminiscence. All animals of the first five cages of each group were kept until the end of the experiment at 20 dpi, or until the individual endpoint of each animal. One further cage per group contained only two directly inoculated hamsters that were euthanized at 4 dpi to study the viral organ tropism. At the start of the experiment (0 dpi), all seven donor hamsters per group were intranasally inoculated with 70 µl of either 10^5.3^ TCID_50_ per animal rMERS-CoV-EMC/2012 (group 1), or 10^5.65^ TCID_50_ per animal rMERS-CoV-SA/D10540/2023 (group 2), distributed equally into each nostril. The exact inoculum titer was calculated by back-titration of the original material on VeroE6 cells (CCLV-RIE 0929). One day post infection, the uninfected contact hamsters were added to the inoculated donor hamsters. All animals were monitored daily for clinical signs and overall conditions until the end of the experiment at 14 dpi. Body weight was monitored daily until the end of the experiment. Additionally, nasal washes were performed daily until 14 dpi or until each respective endpoint of the individual animals. In brief, 200 µL of PBS was administered per nostril of each individual animal and the reflux was collected in a 2 mL Eppendorf-tube. Animals were kept until 14 dpi to study seroconversion. At each individual endpoint, blood was collected from each animal and used for serological analysis by virus neutralization test (VNT_100_). Moreover, organ samples were collected and analyzed via RT-qPCR to investigate viral tissue tropism.

### RNA extraction and RT-qPCR for viral genome detection

100 µL of swab-, nasal washing- or organ samples were extracted via the NucleoMag Vet kit (Macherey-Nagel, Düren, Germany) according to manufacturer’s instructions on a Biosprint 96 platform (Qiagen, Hilden, Germany). Detection of MERS-CoV genome was enabled by using a published PCR-protocol, specific for the upstream E-region of MERS-CoV using the FAM channel ^53^. Primer sequences shows no mismatches for both of the viruses. On HEX channel ß-actin was detected as reference to confirm functional RNA extraction ^54^. The RT-qPCR reaction was prepared using the qScript XLT One-Step RT-qPCR ToughMix (QuantaBio) Kit. All RT-qPCRs were run on a CFX 96 RT-PCR cycler (BioRad, Hercules, CA, USA). Fluorescence values were collected during the annealing phase.

### RNA extraction and RT-qPCR for hDPP4 detection

Total RNA was isolated from tissues of hDPP4 hamsters using PureLink™ RNA Mini Kit (Invitrogen, USA) according to the manufacturer’s instructions. A total of 1 μg RNA was used for cDNA synthesis by reverse transcription using SuperScript™ III First-Strand Synthesis SuperMix (Invitrogen, USA). Quantitative RT–PCR analysis was performed using PrimeTime™ gene Expression Master Mix (IDT, USA) on a LightCycler 96 instrument (Roche, Germany). Primers and probes were listed in Table 1 and synthesized from IDT. Hamster *PPIA* was used as the reference gene. All reactions were performed in triplicate. The RT-qPCR was performed in a 20 μL volume containing 2 μL of cDNA, 10 μl of Master Mix, and 1 μL of qPCR Assay. Thermal cycling parameters were as follows: 95 °C for 3 min; 40 cycles of 95 °C for 15 s, 60 °C for 60 s, and 4 °C for hold.

### Western blotting

Tissues from hDPP4 hamsters were homogenized in RIPA lysis buffer containing protease and phosphatase inhibitor cocktail, and total protein concentrations were determined using the Pierce BCA Protein Assay Kit (Thermo Scientific, USA). Samples were mixed with Laemmli sample buffer, incubated at 70°C for 10 min for protein denaturation. 30 µg of total protein per sample was separated by electrophoresis on a Mini-Protein TGX Gel, and was then transferred onto a PVDF membrane. After blocking with 5% skim milk, the membrane was probed with either anti-hDPP4 (R&D Systems, USA) or anti-β-actin (Santa Cruz, Biotechnology, USA) primary antibodies, followed by HRP-conjugated secondary antibodies. Blots were imaged using a ChemiDoc Imaging System with Clarity Western ECL Substrate Kit (Bio-Rad, USA).

### Titration

Virus titers in selected organ samples were quantified using a TCID_50_ endpoint dilution assay on VeroE6 TMPRSS cells. Briefly, tenfold serial dilutions of each sample were prepared and inoculated in duplicate onto confluent VeroE6 TMPRSS cell monolayers in 96-well plates. Following incubation for 72 h at 37 °C, viral titers were assessed based on the presence of virus-specific cytopathic effects (CPE) and calculated using the measure of infectious dose in specific infections (midSIN) method ^55^.

### Serology

Neutralizing antibodies were determined by performing a virus neutralization assay (VNT). In brief, serum samples were serially diluted from a starting dilution of 1:16 in a 96-well plate and were subsequently mixed with 10^3.3^ TCID_50_/mL of one of the respective MERS-CoV strains used. Following a 1-hour incubation time, 100 µL of VeroE6 cells per well were seeded to the serum-virus mixture containing wells. 72h after incubation at 37 °C, the virus neutralization titer was determined by eye under a light microscope. A well was considered neutralizing in the absence of a MERS-CoV-specific cytopathic effect. The final neutralization titer was calculated based on the last dilution in which no CPE was observed, here shown as VNT_100_ values.

### Pathology

The left lung lobe, brain and nose of animals killed at 4 (n=2 for EMC, n=2 for 10540), and 20 dpi (n=1 for EMC, n=2 for 10540) were immersion-fixed in 10% neutral-buffered formalin for 7 days. Following fixation, the nose was decalcified using Osteosoft™ (Merck KGaA, Germany) for 21 days, cut at 4 transverse sections, decalcified for further 72 hours, and washed for 24 hours in 10% neutral-buffered formalin. The brain was transversally trimmed at 4 levels. The total lung and trimmed tissues were embedded in paraffin. Sections of 3-4 μm thickness were mounted on Super-Frost-Plus-Slides (Fisher Scientific GmbH, Germany), dried for 90 minutes at 60°C and stored at RT until further processing. For viral antigen detection, immunohistochemistry (IHC) was performed on serial consecutive sections as described ^56^. The primary anti-MERS-CoV antibody (Sino Biological #A40068-MM10, 1:1250) was applied overnight at 4°C, the secondary biotinylated goat anti-mouse antibody was applied for 30 minutes at room temperature (Vector Laboratories, Burlingame, CA, USA, 1:200). As controls, consecutive sections were labeled with an irrelevant antibody targeting the capsid protein of Rustrela virus (clone 2H11B1, in house, generated by Sven Reiche^57^.

## Acknowledgment

We gratefully thanks Robin Brandt and Mareen Grawe for their dedicated technical assistance and also the animal caretaker team.

This work was funded by the DURABLE project, co-funded by the European Union, 561 under the EU4Health Programme (EU4H), project no. 101102733.

## References

1 Santacroce, L., Charitos, I. A., Carretta, D. M., De Nitto, E. & Lovero, R. The human coronaviruses (HCoVs) and the molecular mechanisms of SARS-CoV-2 infection. J Mol Med (Berl) 99, 93–106 (2021). 10.1007/s00109-020-02012-8

2 Zaki, A. M., van Boheemen, S., Bestebroer, T. M., Osterhaus, A. D. & Fouchier, R. A. Isolation of a novel coronavirus from a man with pneumonia in Saudi Arabia. N Engl J Med 367, 1814–1820 (2012). 10.1056/NEJMoa1211721

3 WHO. MERS-CoV worldwide overview: Situation update 17 May 2025, <https://data.who.int/dashboards/mers/cases> (2023).

4 Alzahrani, A. et al. Surveillance and Testing for Middle East Respiratory Syndrome Coronavirus, Saudi Arabia, March 2016-March 2019. Emerg Infect Dis 26, 1571–1574 (2020). 10.3201/eid2607.200437

5 Lambrou, A. S. et al. Update on the Epidemiology of Middle East Respiratory Syndrome Coronavirus - Worldwide, 2017-2023. MMWR Morb Mortal Wkly Rep 74, 313–320 (2025). 10.15585/mmwr.mm7419a1

6 Cui, J., Li, F. & Shi, Z. L. Origin and evolution of pathogenic coronaviruses. Nat Rev Microbiol 17, 181–192 (2019). 10.1038/s41579-018-0118-9

7 Han, H. J., Yu, H. & Yu, X. J. Evidence for zoonotic origins of Middle East respiratory syndrome coronavirus. J Gen Virol 97, 274–280 (2016). 10.1099/jgv.0.000342

8 Ithete, N. L. et al. Close relative of human Middle East respiratory syndrome coronavirus in bat, South Africa. Emerg Infect Dis 19, 1697–1699 (2013). 10.3201/eid1910.130946

9 Hassan, A. M. et al. Ongoing Evolution of Middle East Respiratory Syndrome Coronavirus, Saudi Arabia, 2023-2024. Emerg Infect Dis 31, 57–65 (2025). 10.3201/eid3101.241030

10 Khalafalla, A. I. et al. Isolation and genetic characterization of MERS-CoV from dromedary camels in the United Arab Emirates. Front Vet Sci 10, 1182165 (2023). 10.3389/fvets.2023.1182165

11 Kiyong’a, A. N. et al. Middle East Respiratory Syndrome Coronavirus (MERS-CoV) Seropositive Camel Handlers in Kenya. Viruses 12 (2020). 10.3390/v12040396

12 Azhar, E. I. et al. Middle East respiratory syndrome coronavirus-a 10-year (2012-2022) global analysis of human and camel infections, genomic sequences, lineages, and geographical origins. Int J Infect Dis 131, 87–94 (2023). 10.1016/j.ijid.2023.03.046

13 Ogoti, B. et al. Epidemiology and genomic features of MERS coronavirus in Africa: a systematic and meta-analysis review. International Journal of Infectious Diseases, 108456 (2026).

14 Chu, D. K. W. et al. MERS coronaviruses from camels in Africa exhibit region-dependent genetic diversity. Proc Natl Acad Sci U S A 115, 3144–3149 (2018). 10.1073/pnas.1718769115

15 Schroeder, S. et al. Functional comparison of MERS-coronavirus lineages reveals increased replicative fitness of the recombinant lineage 5. Nat Commun 12, 5324 (2021). 10.1038/s41467-021-25519-1

16 Te, N., Rodon, J., Perez, M., Segales, J., Vergara-Alert, J. & Bensaid, A. Enhanced replication fitness of MERS-CoV clade B over clade A strains in camelids explains the dominance of clade B strains in the Arabian Peninsula. Emerg Microbes Infect 11, 260–274 (2022). 10.1080/22221751.2021.2019559

17 Ngere, I. et al. Outbreak of Middle East Respiratory Syndrome Coronavirus in Camels and Probable Spillover Infection to Humans in Kenya. Viruses 14 (2022). 10.3390/v14081743

18 Winstone, H., Stillwell, H., Tan, L. H., Cohen, N. A., Perera, R. A. & Weiss, S. R. Clade C MERS-CoV camel strains vary in protease utilization during viral entry. Proceedings of the National Academy of Sciences 123, e2525313123 (2026).

19 Ogoti, B. M. et al. Biphasic MERS-CoV incidence in nomadic dromedaries with putative transmission to humans, Kenya, 2022–2023. Emerging Infectious Diseases 30, 581 (2024).

20 Mok, C. K. P. et al. T-cell responses to MERS coronavirus infection in people with occupational exposure to dromedary camels in Nigeria: an observational cohort study. The Lancet Infectious Diseases 21, 385–395 (2021).

21 Mok, C. K. et al. Middle East Respiratory Syndrome Coronavirus–Specific T-Cell Responses in Dromedary Camel Abattoir Workers in Nigeria Suggests Frequent Zoonotic Spillover. The Journal of Infectious Diseases, jiag095 (2026).

22 Karani, A. et al. Low-level zoonotic transmission of clade C MERS-CoV in Africa: insights from scoping review and cohort studies in hospital and community settings. Viruses 17, 125 (2025).

23 Bernard-Stoecklin, S. et al. Comparative Analysis of Eleven Healthcare-Associated Outbreaks of Middle East Respiratory Syndrome Coronavirus (Mers-Cov) from 2015 to 2017. Sci Rep 9, 7385 (2019). 10.1038/s41598-019-43586-9

24 Drosten, C. et al. Transmission of MERS-coronavirus in household contacts. N Engl J Med 371, 828–835 (2014). 10.1056/NEJMoa1405858

25. WHO. Middle East respiratory syndrome coronavirus - Kingdom of Saudi Arabia. (2025).

26 Subissi, L. et al. Human MERS-CoV cases are falling but pose an ongoing pandemic threat. Nature Health, 1–4 (2026).

27 de Wit, E. et al. The Middle East respiratory syndrome coronavirus (MERS-CoV) does not replicate in Syrian hamsters. PLoS One 8, e69127 (2013). 10.1371/journal.pone.0069127

28 Raj, V. S. et al. Adenosine deaminase acts as a natural antagonist for dipeptidyl peptidase 4-mediated entry of the Middle East respiratory syndrome coronavirus. J Virol 88, 1834–1838 (2014). 10.1128/JVI.02935-13

29 van Doremalen, N. et al. Host species restriction of Middle East respiratory syndrome coronavirus through its receptor, dipeptidyl peptidase 4. J Virol 88, 9220–9232 (2014). 10.1128/JVI.00676-14

30 van Doremalen, N. & Munster, V. J. Animal models of Middle East respiratory syndrome coronavirus infection. Antiviral Res 122, 28–38 (2015). 10.1016/j.antiviral.2015.07.005

31 Raj, V. S. et al. Dipeptidyl peptidase 4 is a functional receptor for the emerging human coronavirus-EMC. Nature 495, 251–254 (2013).

32 Fan, Z. et al. Efficient gene targeting in golden Syrian hamsters by the CRISPR/Cas9 system. PloS one 9, e109755 (2014).

33 Li, R. et al. Production of genetically engineered golden Syrian hamsters by pronuclear injection of the CRISPR/Cas9 complex. Journal of visualized experiments: JoVE, 56263 (2018).

34 Gibson, S. A. et al. Differences in susceptibility to SARS-CoV-2 infection among transgenic hACE2-hamster founder lines. Viruses 16, 1625 (2024).

35 van Doremalen, N. et al. Host species restriction of Middle East respiratory syndrome coronavirus through its receptor, dipeptidyl peptidase 4. Journal of virology 88, 9220–9232 (2014).

36 Zhao, J. et al. Rapid generation of a mouse model for Middle East respiratory syndrome. Proceedings of the National Academy of Sciences 111, 4970–4975 (2014).

37 Agrawal, A. S. et al. Generation of a transgenic mouse model of Middle East respiratory syndrome coronavirus infection and disease. Journal of virology 89, 3659–3670 (2015).

38 Li, K. et al. Middle East respiratory syndrome coronavirus causes multiple organ damage and lethal disease in mice transgenic for human dipeptidyl peptidase 4. The Journal of infectious diseases 213, 712–722 (2016).

39 Tao, X. et al. Characterization and demonstration of the value of a lethal mouse model of Middle East respiratory syndrome coronavirus infection and disease. Journal of virology 90, 57–67 (2016).

40 van Doremalen, N. et al. Transmission dynamics of MERS-CoV in a transgenic human DPP4 mouse model. npj Viruses 2, 36 (2024).

41 Shi, L. et al. A single-dose intranasal immunization with a novel bat influenza A virus-vectored MERS vaccine provides effective protection against lethal MERS-CoV challenge. mBio, e01107–01125 (2025).

42 Arabi, Y. M. et al. Clinical course and outcomes of critically ill patients with Middle East respiratory syndrome coronavirus infection. Annals of internal medicine 160, 389–397 (2014).

43 Halfmann, P. J. et al. SARS-CoV-2 Omicron virus causes attenuated disease in mice and hamsters. Nature 603, 687–692 (2022).

44 Uraki, R. et al. Characterization of SARS-CoV-2 Omicron BA. 4 and BA. 5 isolates in rodents. Nature 612, 540–545 (2022).

45 Golden, J. W. et al. Hamsters expressing human angiotensin-converting enzyme 2 develop severe disease following exposure to SARS-CoV-2. MBio 13, e02906–02921 (2022).

46 Cong, Y. et al. Longitudinal analyses using 18F-Fluorodeoxyglucose positron emission tomography with computed tomography as a measure of COVID-19 severity in the aged, young, and humanized ACE2 SARS-CoV-2 hamster models. Antiviral Research 214, 105605 (2023).

47. Gilliland, T., et al. Transchromosomic bovine-derived anti-SARS-CoV-2 polyclonal human antibodies protects hACE2 transgenic hamsters against multiple variants. Iscience 26 (2023).

48 Oladunni, F. S. et al. Lethality of SARS-CoV-2 infection in K18 human angiotensin-converting enzyme 2 transgenic mice. Nature communications 11, 6122 (2020).

49 Rathnasinghe, R. et al. Comparison of transgenic and adenovirus hACE2 mouse models for SARS-CoV-2 infection. Emerging microbes & infections 9, 2433–2445 (2020).

50 Park, J.-E. et al. Proteolytic processing of Middle East respiratory syndrome coronavirus spikes expands virus tropism. Proceedings of the National Academy of Sciences 113, 12262–12267 (2016).

51 Muth, D. et al. Transgene expression in the genome of Middle East respiratory syndrome coronavirus based on a novel reverse genetics system utilizing Red-mediated recombination cloning. Journal of General Virology 98, 2461–2469 (2017).

52 Thi Nhu Thao, T., et al. Rapid reconstruction of SARS-CoV-2 using a synthetic genomics platform. Nature 582, 561–565 (2020).

53 Coleman, C. M. & Frieman, M. B. Growth and quantification of MERS-CoV infection. Current protocols in microbiology 37, 15E. 12.11–15E. 12.19 (2015).

54 Wernike, K., Hoffmann, B., Kalthoff, D., König, P. & Beer, M. Development and validation of a triplex real-time PCR assay for the rapid detection and differentiation of wild-type and glycoprotein E-deleted vaccine strains of Bovine herpesvirus type 1. Journal of virological methods 174, 77–84 (2011).

55 Cresta, D. et al. Time to revisit the endpoint dilution assay and to replace the TCID50 as a measure of a virus sample’s infection concentration. PLoS computational biology 17, e1009480 (2021).

56 Britzke, T. et al. Live attenuated SARS-CoV-2 vaccine OTS-228 demonstrates efficacy, safety, and stability in preclinical model. npj Vaccines 10, 104 (2025).

57 Matiasek, K. et al. Mystery of fatal ‘staggering disease’unravelled: novel rustrela virus causes severe meningoencephalomyelitis in domestic cats. Nature Communications 14, 624 (2023).

