## Supplement Figure 1 for "First evaluation of a human DPP4 transgenic hamster model for MERS-CoV pathogenesis and transmission"

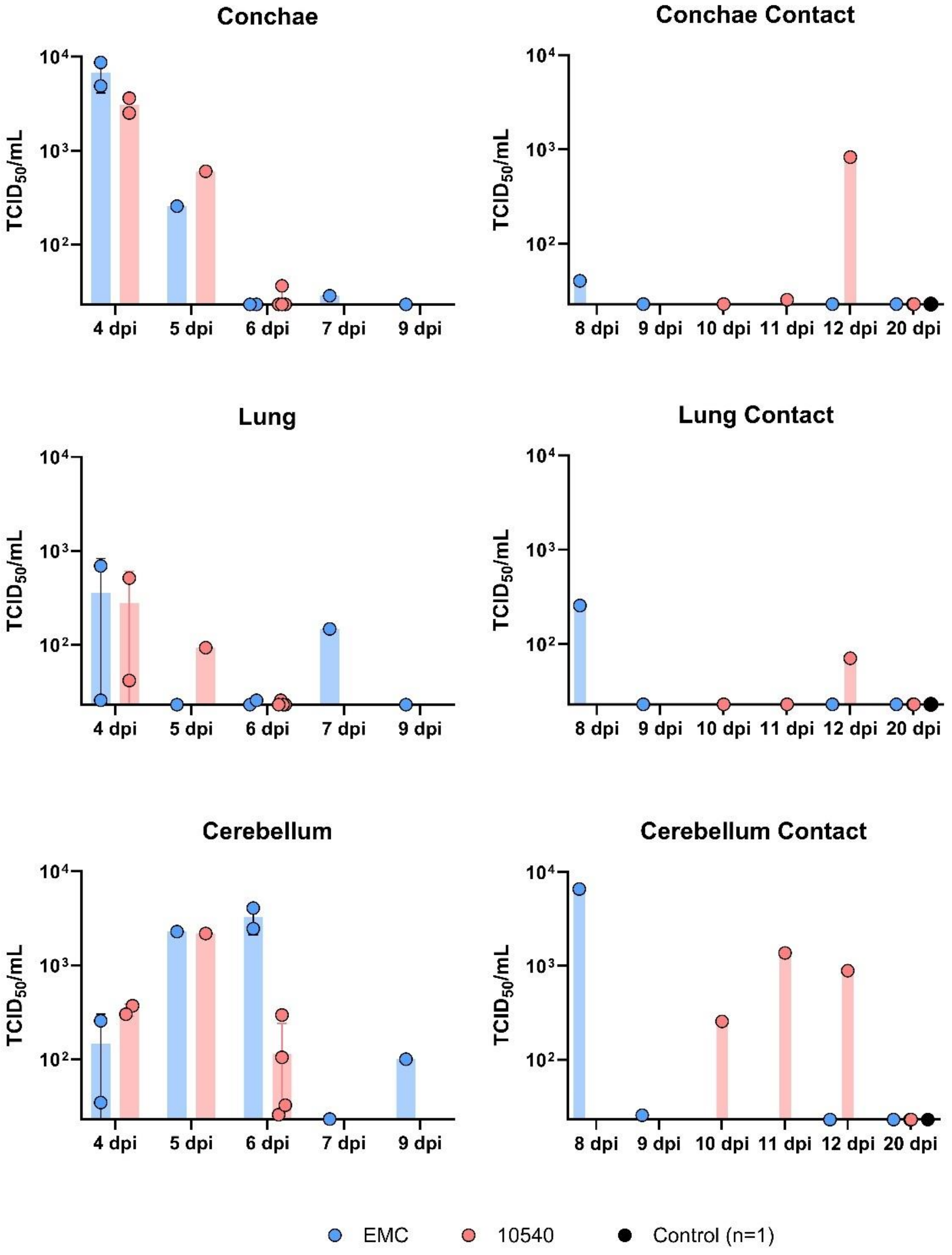


**Supplementary Figure 1: Infectious MERS-CoV ($TCID_{50}$/mL) in organ samples.** Virus titers in selected organs (Conchae, Lung, Cerebellum) were determined by endpoint serial dilution assay on VeroE6 TMPRSS cells. TCID50/mL titers were assessed after 72 h based on cytopathic effects (CPE) and calculated by the midSIN method.
