## Supplement Figure 2 for "First evaluation of a human DPP4 transgenic hamster model for MERS-CoV pathogenesis and transmission"

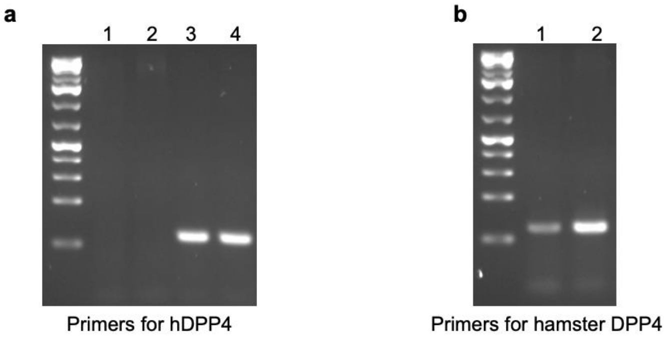


**Supplementary Figure 2. PCR validation of hDPP4- and hamster DPP4-specific primers.** (a) PCR products amplified with hDPP4-specific primers were sequenced and confirmed as hDPP4. Lanes 1–4 represent cDNA from wild-type hamster lung, wild-type hamster spleen, hDPP4 transgenic hamster lung, and hDPP4 transgenic hamster spleen, respectively. (b) PCR products amplified with hamster DPP4-specific primers were sequenced and confirmed as hamster DPP4. Lanes 1 and 2 represent cDNA from hDPP4 transgenic hamster lung and spleen, respectively.
