## Supplement Table 1 for "First evaluation of a human DPP4 transgenic hamster model for MERS-CoV pathogenesis and transmission"

**Supplementary Table 1: Clinical picture and necropsy timepoint.**

| virus group | dpi | animal ID | cage ID | function | sex | clinical manifestation |
|---|---|---|---|---|---|---|
| **EMC** | d7 | 1 | 1 | Donor | m | unsteady gait / ataxic gait, difficulty grasping, head tilt |
| **EMC** | d8 | 2 | 1 | Contact | m | apathetic, bloated/distended abdomen |
| **EMC** | d6 | 3 | 2 | Donor | m | found dead |
| **EMC** | d9 | 4 | 2 | Contact | m | found dead |
| **EMC** | d6 | 5 | 3 | Donor | m | unsteady, collapses |
| **EMC** | d20 | 6 | 3 | Contact | m | scheduled euthanasia |
| **EMC** | d6 | 7 | 4 | Donor | f | unsteady, bloated/distended abdomen |
| **EMC** | d12 | 8 | 4 | Contact | f | neurological symptoms, head tilt, unsteady/ataxic gait |
| **EMC** | d6 | 9 | 5 | Donor | f | unsteady / ataxic |
| **EMC** | d12 | 10 | 5 | Contact | f | neurological symptoms, head tilt, unsteady/ataxic gait |
| **EMC** | d4 | 11 | 6 | Donor | f | scheduled euthanasia |
| **EMC** | d4 | 12 | 7 | Donor | m | scheduled euthanasia |
| **10540** | d6 | 13 | 8 | Donor | m | unsteady / ataxic |
| **10540** | d10 | 14 | 8 | Contact | m | head tilt, unsteady |
| **10540** | d6 | 15 | 9 | Donor | m | unsteady / ataxic |
| **10540** | d12 | 16 | 9 | Contact | m | *severely apathetic* |
| **10540** | d5 | 17 | 10 | Donor | m | severe dyspnea |
| **10540** | d10 | 18 | 10 | Contact | m | pronounced head tilt, unsteady |
| **10540** | d6 | 19 | 11 | Donor | f | unsteady / ataxic |
| **10540** | d20 | 20 | 11 | Contact | f | scheduled euthanasia |
| **10540** | d6 | 21 | 12 | Donor | f | unsteady / ataxic |
| **10540** | d20 | 22 | 12 | Contact | f | scheduled euthanasia |
| **10540** | d4 | 23 | 13 | Donor | f | scheduled euthanasia |
| **10540** | d4 | 24 | 14 | Donor | m | scheduled euthanasia |
| **10540** | d20 | 25 | 15 | neg. Control | m | scheduled euthanasia |
